# A taxonomic synopsis of *Heterotis* (Melastomataceae), including *H. kupensis*, a new threatened species endemic to Mt Kupe, Cameroon

**DOI:** 10.1101/2025.08.26.672404

**Authors:** Vida J. Svahnström, Martin Cheek

## Abstract

We provide a synopsis of the African genus *Heterotis* (Melastomataceae), recognising seven species and including the description of a new species to science *Heterotis kupensis* Sv. & Cheek (Melastomataceae) which is only known from Mt Kupe in the South West Region of Cameroon. In addition, we select a lectotype for *H. prostrata* (Thonn.) Benth., second step lectotype for *H. buettneriana* (Cogn. ex Büttner) Jacq.-Fél. and create the new combination *H. welwitschii* (Cogn.) Sv. & Cheek to replace *H. cogniauxiana* (A.Fern. & R.Fern.) Ver.-Lib. & G.Kadereit synon. nov. Distribution, habitat, phenology, taxonomic notes, and preliminary conservation assessments following IUCN Red List criteria are presented along with a key to the species.

## Introduction

Melastomateae is a morphologically diverse and species-rich pantropical tribe of Melastomataceae characterised by cochleate seeds with tubercles, a crown of bristles on the apex of the ovary and staminal pedoconnectives with bifurcated ventral appendages (Veranso-Libalah et al. 2022). The Melastomateae is the most species-rich melastome tribe in Africa, consisting of 24 genera and about 143 species (Veranso-Libalah et al. 2020, Chen et al. 2025). Delimitation of genera in African Melastomateae has historically been challenging due to extensive diversity and variation in a multitude of morphological characters, exacerbated by the apparent prevalence of evolutionarily labile traits within the tribe. The genus *Heterotis* Benth. was first recognised by Bentham (1849) in the earliest treatment of African Melastomateae, who proposed four sections *Heterotis* Benth., *Cyclostemma* Benth., *Leiocalyx* Benth., and *Wedeliopsis* Benth. In his treatment, Bentham (1849) defined *Heterotis* on the basis of its persistent, laciniate calyces with setose appendages on the hypanthium, dimorphic anthers, and dehiscent loculicidal capsules. Triana (1872) transferred all of the species in sections *Leiocalyx* and *Wedeliopsis* to *Tristemma* Juss. and sections *Heterotis* and *Cyclostemma* were subsumed into a broadly defined *Dissotis* as sect. *Heterotis*, which was elevated to subgenus *Heterotis* (Benth.) A.Fern. & R.Fern by Fernandes & Fernandes (1969). Jacques-Félix (1981) resurrected *Heterotis* at genus rank, recognising that staminal dimorphism, the defining character of Triana (1872)’s concept for *Dissotis*, is an unreliable trait for generic classification. He defined three sections within *Heterotis*: sect. *Heterotis*, sect. *Argyrella* (Naudin) Jacq.-Fél. and sect. *Cyclostemma*.

Recently, the African Melastomateae has undergone revision in light of phylogenetic study, resulting in an updated classification of 24 monophyletic genera each diagnosed by combinations of morphological characters and supported by generic keys (Veranso-Libalah et al. 2017, Veranso-Libalah et al. 2020, Chen et al. 2025). Many accepted genera were found to be polyphyletic, including *Heterotis.* Traits which have previously been used to define genera in this tribe such as flower merosity, habit, and isomorphic vs. dimorphic stamens are now, in a phylogenetic framework, recognized to have been gained and lost many times. As a result, *Heterotis* was recircumscribed, resulting in a more narrowly defined, monophyletic genus, consisting only of section *Heterotis* sensu Jacques-Félix (1981).

Veranso-Libalah et al. (2017) distinguish *Heterotis* morphologically from other African Melastomateae on the basis of the following combination of characters: decumbent herbs, leaves orbicular to ovate-lanceolate, hypanthium with stalked emergences, flowers large, 5-merous, calyx-lobes persistent, apex of intersepalar appendages with stellate emergences, and seeds cochleate and visibly arillate. Six species are currently recognised in the genus *Heterotis*, three of which are restricted to the Guinean-Congolian region (*H. fruticosa* (Brenan) Ver.-Lib. & G.Kadereit, *H. cogniauxiana* (A.Fern. & R.Fern.) Ver.-Lib. & G.Kadereit, and *H. buettneriana* (Cogn. ex Büttner) Jacq.-Fél.), whereas the remaining three are more widely distributed in tropical Africa (*H. decumbens* (P.Beauv.) Jacq.-Fél., *H. prostrata* (Thonn.) Benth., and *H. rotundifolia* (Sm.) Jacq.-Fél.). Within species of the genus, several characters exhibit extensive variation, including some of which have previously been used for species delimitation, such as hypanthium indumentum, leaf shape, and habit. As a result, there has been confusion in distinguishing between species. Previous treatments have considered *H. prostrata* as a synonym (Wickens 1975) or variety of *H. rotundifolia* (Jacques-Félix 1971) and *H. rotundifolia* as indistinguishable from *H. decumbens* (Gilbert 1995). *H. fruticosa* was originally described as a variety of *H. rotundifolia* before being elevated to species level (Brenan 1950, Keay 1952). In the most recent species-level treatment of the genus, Jacques-Félix (1981, 1995) recognised six species but suggested that they are closely related, and that the widespread and particularly problematic and variable *H. rotundifolia*, *H. prostata* and *H. decumbens* could represent a species complex.

No new species have been added to *Heterotis* s.s. since the description of *H. fruticosa* (as *Dissotis rotundifolia var. fruticosa*) (Brenan 1950). During fieldwork in 1995 on Mt. Kupe in Cameroon, a distinctive species of Melastomateae was collected from grassland at the summit. It was first recognised as a provisionally new taxon as *Dissotis* sp. 1 in a checklist for Mt. Kupe (Cheek et al. 2004). The generic keys and descriptions in Veranso-Libalah (2017, 2020) support this species placement in the genus *Heterotis* based on the presence of stalked stellate emergences on the hypanthium, intersepalar appendages with stellate hairs at the apex, cymose inflorescences, and large, 5-merous flowers.

Here, we provide a taxonomic synopsis of the genus *Heterotis*, including a new combination, a lectotypification and a second step lectotypification, and a description of a previously unnamed species from Cameroon. We also present preliminary conservation assessments and a key to all the species in the genus.

**Figure 1:**
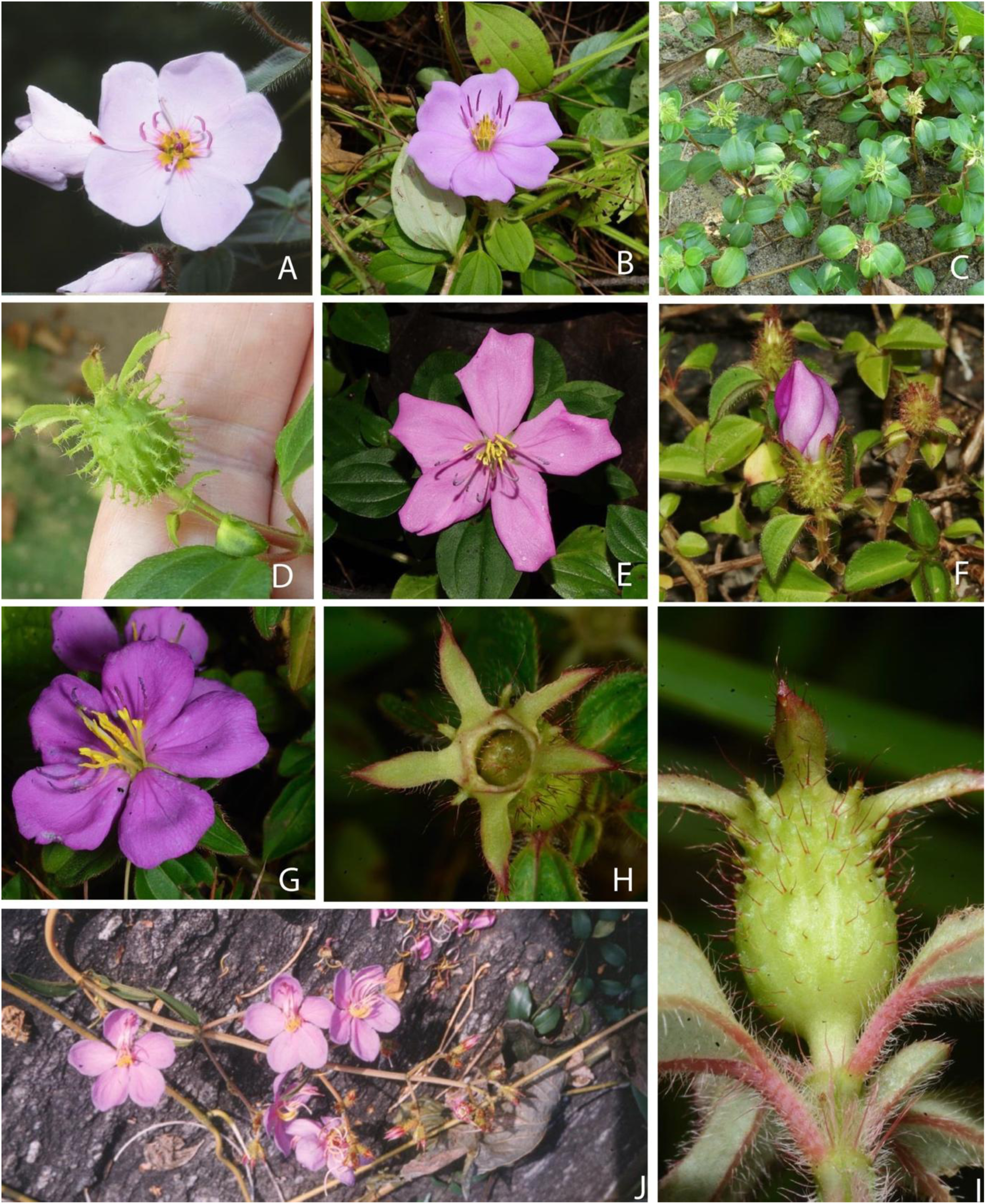
Representatives of the genus *Heterotis*. **A** *Heterotis kupensis* sp. nov.; **B-D** *H. prostrata* flower (B), habit (C) and hypanthium and calyx (D)**; E-F** *H. rotundifolia* flower (E) and plant (F); **G-I** *H. decumbens* flower (G), calyx (H) and hypanthium (I); **J** *H. fruticosa*. Photos: **A** Martin Cheek; **B** Hubert Lagrange; **C-D** Carel Jongkind; **E** Vida J. Svahnström; **F** James Bailey; **G-I** Ehoarn Bidault; **J** RBG Kew.

## Materials and Methods

This study is predominantly based on herbarium specimens held at K, and in some cases images from other herbaria including BR, BM, COI, LISU, G-DC, P, NY, US, WAG, MO, G, and C. Herbarium codes follow Thiers (continuously updated). The description of the new species is based on seven duplicates of the type, which comprise all of the known material for this taxon at present, collected according to Cheek & Cable (1997). Material of the new species was compared morphologically with material of all other species of *Heterotis* held at K. All of the specimens seen by the authors are indicated “!” whereas specimens for which we examined the images are marked ‘image’. Images (GBIF, continuously updated) were usually of insufficient resolution to enable or confirm identifications. There was insufficient time to obtain material on loan to K for this study and no funds available for herbarium visits for relation to this work. Herbarium material was examined using a Leica M165C dissecting binocular microscope. The drawing was made using a Leica Wild M8 dissecting binocular microscope fitted with an eyepiece graticule measuring in units of 0.025 mm at maximum magnification with a Leica 308700 camera lucida attachment. Type specimens are not given for all homotypic names in synonymy. The conservation assessment follows the categories and criteria of the International Union for the Conservation of Nature (IUCN, 2012) and the guidelines for their use (IUCN 2024).

### Taxonomic Treatment

**Heterotis** *Benth.* (Bentham 1849: 347). Type species: *Heterotis rotundifolia* (Sm.) Jacq.-Fél.

### Key to the species of *Heterotis*

#### 1. Heterotis kupensis *Sv. & Cheek* sp. nov. (Fig. 2 & 3)

Type: Cameroon, S.W. Province, NE side of Mt Kupe, Peak No. 2, 1900 m elev., fl. 11 Jan. 1995, *Sidwell* K. 428 with *Cheek*, *Gosline*, *Okah*, *Etuge*, *Oyugi* (holotype K; isotypes BR, CAS, K, SCA, YA) *Dissotis sp. 1* Cheek & Woodgyer in Cheek et al. (2004: 335 & plate 9F) *Erect herb* 55 – 75 cm tall, with multiple stems from the base, stems ascending, woody at the base, unbranched their entire length, or sparingly and shortly branched in the distal half, sometimes rooting at nodes, stems with internodes 3.2 – 6.5 cm long, becoming shorter up the stem, covered with spreading hairs (1.0 – ) 1.5 – 4.5 ( – 5.5) mm long, scattered on internodes and in a dense ring at nodes. *Leaves* simple, opposite, elliptic or elliptic-ovate, drying brown, leaves (0.8 – ) 2.4 – 3.6 x (0.5 – ) 1.2 – 1.9 cm, decreasing in size up the stem, each leaf with apex acute, base rounded, margins entire to minutely crenulate with appressed bristle bases along entire length, bristles 1.5 –3.0 mm long; midnerve and two pairs of lateral nerves impressed on adaxial surface of lamina, prominent on abaxial surface, lateral nerves diverging from the midnerve at the base of the blade, curvilinear, outer pair of lateral nerves terminating at leaf margin 3/4 of the length of the lamina, inner pair of lateral nerves meeting or nearly meeting the midnerve at the leaf apex, transverse veins only faintly visible, forming weak arches. Adaxial surface of lamina covered (20% cover) with simple antrorse strigose hairs, more or less sigmoid, 1.0 – 4.5 mm with a swollen botuliform base appressed ca. 0.5 mm long, the distal part fine, bristlelike, hairs becoming shorter towards the apex of lamina. Abaxial surface (10% cover) with simple antrorse, slightly appressed hairs, with less pronounced swollen bases than hairs on adaxial surface, 1.0 – 3.8 mm long, hairs sparse on abaxial lamina surface, denser on the principal veins; petiole terete 2.0 – 3.5 mm long, hairs as lamina. *Inflorescence* terminal, cymose, (1–) 4 – 5 (– 6)-flowered, or with 1– 2 flowers in upper axils, flowers 5-merous, pedicels 2 – 4 mm long. *Bracts* caducous, falling at anthesis, two pairs per flower, the outer pair 3.0 – 4.5 x 2.0 mm drying tan, ovate, glabrous except for ciliate margins with cilia 1.0-3.5 mm long, inserted 1.5 – 2.0 mm above the most distal leaf node, inner pair inserted 2.0 mm above the outer pair and 1.0 mm below the flower, drying purple, ovate, 2.0 – 2.7 x 1.5 – 2.5 mm, glabrous except for ciliate margins with cilia 0.8 – 1.3 mm long. *Hypanthium* campanulate 5.0 – 9.5 x 4.3 – 6.0 mm, densely covered with persistent white hairs 2.5– 3.5 mm long with swollen bases, penicillate appendages on the upper ½ of hypanthium 0.3 – 0.9 x 0.2 – 0.4 mm with a tuft of 1 – 4 white hairs, upper ½ of hypanthium also with purple, filiform appendages 1.2 – 2.0 x 0.2 – 0.3 mm, stellate with many white hairs at apex; intersepalar appendages similar in form and colour to the filiform hypanthial appendages but caducous, larger, 2.0 – 2.9 x 0.2 – 0.4 ( – 0.8) mm. *Calyx* green drying purple, sepals oblong, 5.0 – 7.5 x 2.5 – 3.5 mm, tardily caducous, falling after anthesis, margins ciliate with cilia (0.1 – ) 0.5 – 1.0 ( – 2.0) mm long, glabrous on the inside, with very few short hairs near base on the outside 0.4 – 1.4 mm long, and a thickened stellate cluster of many white hairs each 1.0 – 4.0 mm long at the apex. *Corolla* pale pink, centre yellow with a dark pink edge, petals five, obovate, 2.0 – 2.5 x 2.0 – 2.5 cm, glabrous except for ciliate margins with cilia 0.2 – 0.5 ( – 1.5) mm long. *Stamens* ten, yellow with pink anthers opening by an introrse pore, dimorphic, antesepalous stamens five, filament 3.0 – 4.0 x 0.5 – 1.0 mm, pedoconnective hastate, 3.5 – 4.5 mm long,1.0 – 1.5 mm wide at base, tapering at the apex, the proximal arms approximately triangular, diverging from the truncate base of the connective and each other by c. 180**°,** ventral appendage pair swellings transversely ellipsoid, 0.5 – 0.7 mm collateral at the connective base, anther falcate, 4.5 – 5.0 x 0.5 – 1.0 mm, antepetalous stamens five, filaments 2.5 – 3.5 x 0.5 – 1.0 mm, pedoconnective not developed, ventral appendages spreading, each ovoid 0.2 – 0.3 mm long, diverging from each other by 90°, anther 2.5 – 3.0 x 0.5 – 1.0 mm, erect. *Ovary* ellipsoid ca. 2.0 x 1.5 mm, sparsely covered in white hairs 0.5 – 1.0 mm long, with a ring of five appendages at apex surrounding the style, persistent in fruit, appendages comprised of 5-7 erect bifurcating bristles each 1.0 –1.5 mm long. *Style* ca. 15 mm long, glabrous, curved, stigma truncate. *Fruit* not seen.

**Figure 2:**
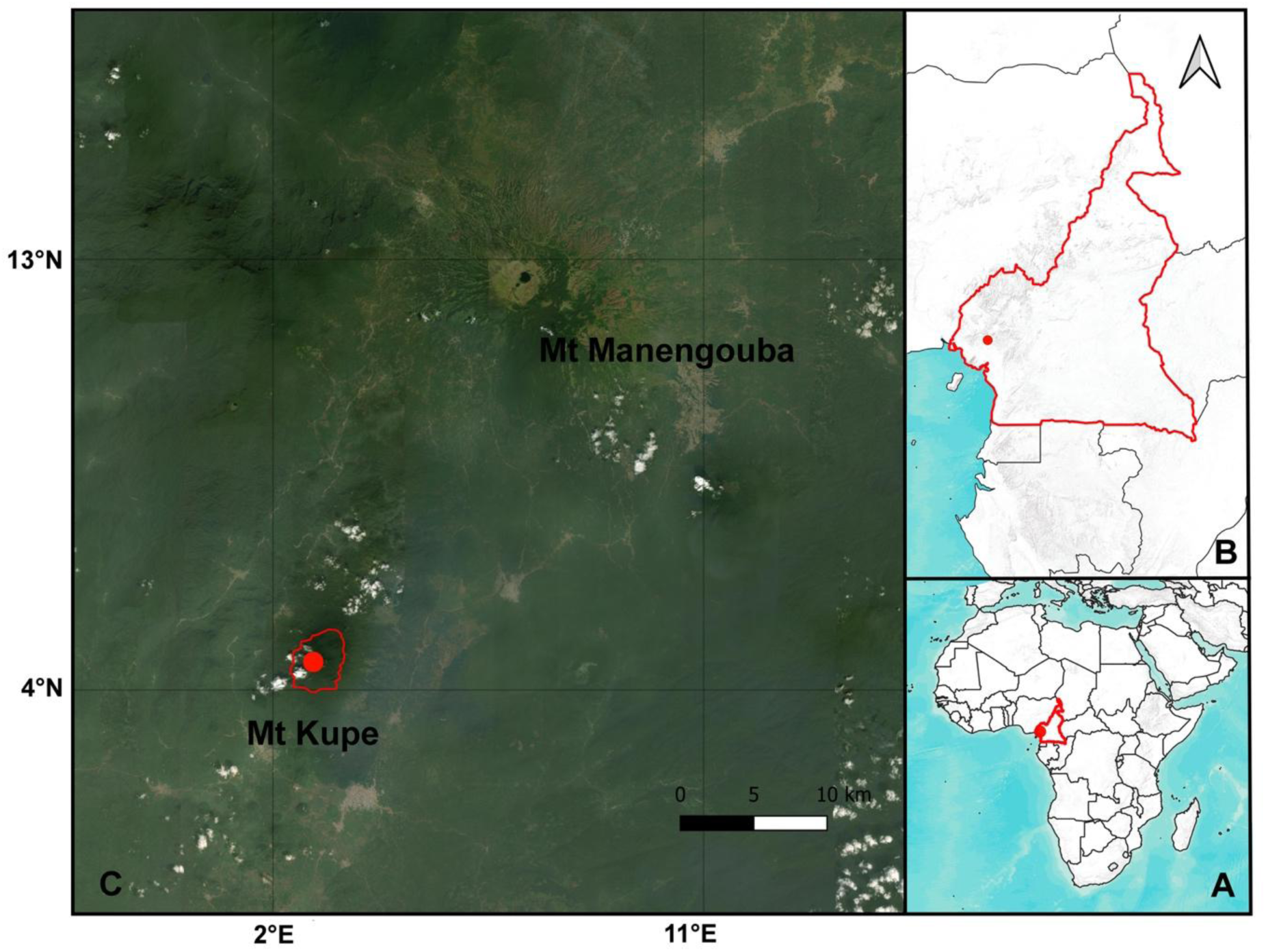
Global distribution map of *Heterotis kupensis*.

**Fig. 3.**
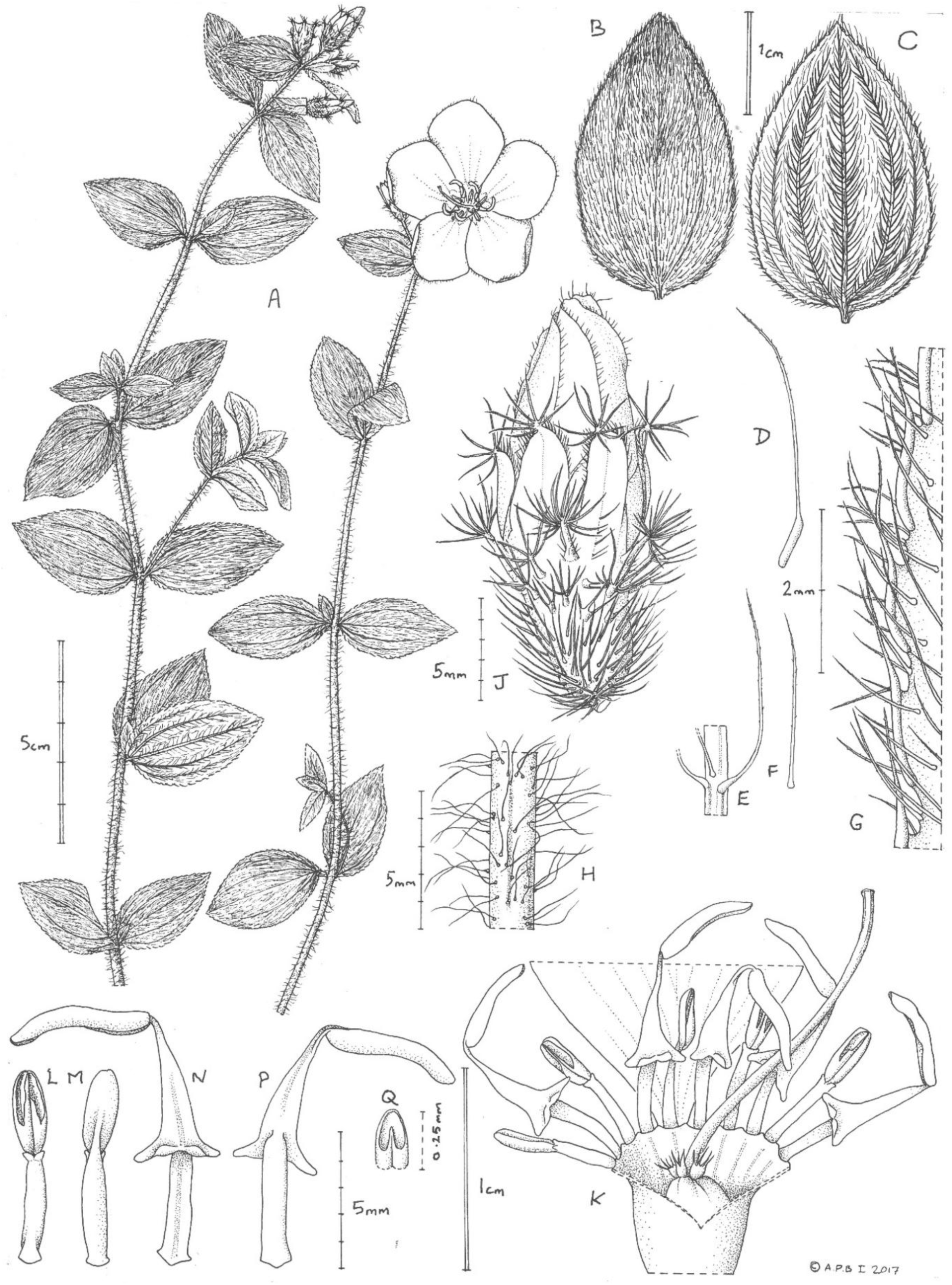
Heterotis kupensis. A. habit (open flower from photo); B. leaf, adaxial surface; C. leaf, abaxial surface; D. hair, from leaf lamina, proximal part, adaxial surface; E. hair, from leaf midrib, abaxial surface; F. hair, from leaf lamina, abaxial surface; G. margin of leaf with 2 hair types, abaxial surface; H. stem indumentum; J. hypanthial ornamentation (flower bud); K. hypanthium (hairs omitted) part opened to show ovary apex and stamens, all but one petal removed; L. antepetalous (short) stamen, ventral face; M. as L, dorsal face; N. antesepalous (large) stamen, ventral face; P. as N, dorsal face; Q. detail of N (pore at anther apex). A-Q all from *Sidwell* 428. Drawn by Andrew Brown.

##### Recognition (Diagnosis)

*Heterotis kupensis* differs from all other members of the genus by its tardily caducous, oblong calyx lobes and hastate pedoconnectives on the antesepalous stamens with transversely ellipsoid ventral appendages. All other species of *Heterotis* have antesepalous stamens which lack a hastate base and which have bilobed ventral appendages. Additionally, the new species is unique in that the hypanthium is covered by hairs with thickened bases as well as filiform appendages on the upper half only, whereas the hypanthium is uniformly covered with filiform appendages in all other species, except for *H. decumbens* which has only simple hairs on the hypanthium. The leaves of *H. kupensis* appear subsessile with short (< 3.5 mm) petioles, whereas most specimens of the remaining six species of *Heterotis* are distinctly petiolate. *H. kupensis* is the only species in the genus which has a corolla which a yellow centre; the other species have a uniformly pink (occasionally white) corolla.

##### Distribution and habitat

Cameroon, South West Region. Known only from the summit of Mount Kupe. Montane grassland with *Adenocarpus mannii* and *Pentas sp. c.* 1900 m asl.

##### Preliminary conservation status

This species is only known from the type specimen, collected from montane grassland at the summit of one of the peaks of Mt Kupe. The area of grassland at the peak is less than 1 km^2^, and as such the extent of occurrence (EOO) and area of occupancy (AOO) are both calculated as 4 km^2^ (using 2×2 km grid cells, see IUCN guidelines), well below the thresholds for threatened categories under subcriteria B1 and B2. There are several other peaks on Mt Kupe with grassland areas at the summit; these have been explored by botanists and this species has not been found at any other localities. However, steep, inaccessible areas on these peaks have not been thoroughly surveyed (Cheek *et al*. 2004) and this species may occur on one or more peaks hitherto undetected. Even if this species occurs on these peaks, or in other areas of montane grassland adjacent to Mt Kupe, it is highly likely that the true AOO falls below the threshold for a threatened category, considering the small area of these summit grasslands. Moreover, extensive surveys in other upland areas of the Cameroon Highlands have been carried out and published but this species has so far been found in none of these (Cheek et al. 1996; Cable & Cheek 1998; Cheek et al. 2000; Maisels et al. 2000; Chapman & Chapman 2001; Cheek et al. 2004; Harvey et al. 2004; Cheek et al. 2006; Cheek et al. 2010; Harvey et al. 2010; Cheek et al. 2011). While the population size has not been assessed, and although this species was reported as common at its only known site, it is likely that less than 1,000 individuals occur there given the extremely small area of potentially suitable habitat. There are no known threats to this species at its type locality, and human encroachment does not currently extend beyond ca. 1,100 m asl on Mt Kupe apart from on the eastern slopes. There is a potential future threat of fire, however no evidence of human-set fires has been detected on any of the peaks at Mt. Kupe (Cheek *et al*. 2004). Given the likelihood that this species has an extremely small population size of less than 1,000 individuals, it is preliminarily assessed as Vulnerable D1.

##### Specimens examined

**CAMEROON.** Cameroon, S.W. Province (now Region), NE side of Mt Kupe, Peak No. 2, 1900 m alt, fl. 11 Jan. 1995, *Sidwell* K. 428 with *Cheek*, *Gosline*, *Okah*, *Etuge*, *Oyugi* (holotype K; isotypes BR, CAS, K, SCA, YA)

##### Phenology

Collected in flower in Jan.

##### Etymology

The species is named for Cameroon’s Mt Kupe where it was discovered and to which it currently appears to be endemic. Other species endemic to and named for Mt Kupe include *Memecylon kupeanum* R.D. Stone *et al*. (Melastomataceae, Stone *et al*. 2008), *Afrothismia kupensis* Cheek (Afrothismiaceae, Cheek *et al*. 2019, 2024) and *Kupeantha kupensis* Cheek & Sonké (Rubiaceae, Cheek *et al*. 2018).

##### Notes

Each of the three peaks of Mt Kupe have small areas of montane grassland with similar plant assemblages (Cheek *et al*. 2004), yet *H. kupensis* has only been collected from one of these, where it was recorded as being locally common. These areas of montane grassland are considered to be inselberg caps of their respective peaks (Cheek *et al*. 2004). Two other species of *Heterotis* (*H. fruticosa* and *H. cogniauxiana*) occur on inselbergs in Nigera and Angola respectively. These are also the only other species in the genus which consistently exhibit an upright, woody growth habit, whereas the remaining species are typically decumbent herbs that root at the nodes.

#### 2. Heterotis welwitschii *(Cogn.) Sv. & Cheek* comb. nov

Basionym: *Osbeckia welwitschii* Cogn. (Cogniaux 1891: 333) Type: Angola, Pungo Andongo, rio Cuanza, pr. de Sansamanda, *Welwitsch 907* (holotype G-DC barcode G00319429! image; isotypes: BM barcode BM000902297! image, COI barcode COI00005417! image, K barcode K000313363!, LISU barcode LISU209417, P barcode P000412573! image) *Dissotis cogniauxiana* A.Fern. & R.Fern. (Fernandes & Fernandes 1954: 67) *Heterotis cogniauxiana* (A.Fern & R.Fern.) Veranso-Libalah & G.Kadereit **synon. nov.**

##### Distribution and habitat

Angola. Growing among rocks on the banks of streams in rocky deciduous woodland, 1,110 –1,300 m elev.

##### Representative specimens examined

**ANGOLA.** Mt. Tchivira. Lucondo Mission near NE corner of mountain, 1110 m elev., fl. 7 Oct. 2013, *Goyder et al.* 7373 (K!); Vila Arriaga, Mahita, fl. & fr. 25 Jan. 1962, *Correia 1798* (LISC barcode LISC030602! image); Cuanza Norte, Pungo Andongo, 1150 m elev., fl. 31 March 1937, *Exell & Mendonça 211* (LISC barcode LISC030603! image)

##### Conservation status

*H. welwitschii* is endemic to the Angolan highlands. Based on herbarium specimens, this species has a minimum EOO of 10,080 km^2^, within the threshold for the Vulnerable category. Plants of Angola (Figueiredo & Smith 2008) lists this species from four provinces: Huila, Namibe, Malange, and Cuanza Norte. It is only represented from a single collecting locality in each of these provinces corresponding to four widely separated locations. However, it is likely that it is more widespread than can be inferred from specimen records due to low botanical collection effort across much of its large range, which extends from the northern to southern Angolan highlands. It is likely undergoing a continuing decline in the area, extent, and suitability of habitat at three of its known localities which are near towns (e.g. Bibala) or in areas which have been extensively converted for agriculture. However, there are large areas of potentially suitable habitat within its extensive range and the most recent collection is from Mt. Tchivira, where the native vegetation is intact. This species could potentially qualify as Vulnerable if it is in fact known from fewer than ten locations undergoing a continuing decline, or it may be Least Concern if found to be more widespread. Thus, given the uncertainty, this species is preliminarily assessed as Near Threatened B1b(iii).

##### Phenology

Collected in flower between Oct. – March, and in fruit Jan. – March.

##### Etymology

Named for the Austrian botanical collector Friedrich Welwitsch (1806 – 1872), who collected the type specimen from which the species was described.

##### Notes

*Osbeckia welwitschii* Cogn. was published in 1891, in the same work as *Dissotis welwitschii* Cogn. (Cogniaux 1891). Thus, when this species was later transfered to *Dissotis* Fernandez and Fernandez (1954) needed a nomen novum: *D. cogniauxiana*. Following its recent placement in *Heterotis* (Veranso-Libalah et al. 2017), the combination *H. cogniauxiana* (A.Fern. & R.Fern) Veranso-Libalah & G.Kadereit was made. However, under the Code (Turland *et al*. 2018) the correct combination is *H. welwitschii*.

#### 3. Heterotis fruticosa *(Brenan) Veranso-Libalah & G.Kadereit* (2017: 609)

Type: Nigeria, Ondo, about 3 km. E of Ondo, north of the Akure road, fl. 21 Sept. 1948 *Keay* 22569 (K barcode K000313153!).

Basionym*: Dissotis rotundifolia var. fruticosa* Brenan (Brenan 1950: 227)

*Dissotis fruticosa* (Brenan) Brenan & Keay (Keay 1953: 547)

##### Distribution and habitat

Idanre Hills, Ondo province in western Nigeria. “Shrubby vegetation among *Trilepis* (now *Afrotrilepis*) mats”. 450 – 950 m elev.

##### Representative specimens examined

**NIGERIA.** Ondo, Idanre Hills, Carter Peak, fl. 24 Oct. 1948, *Keay* 22583 (K barcode K002833085!); Ondo, A few yards to the U.H.F on a wet area on top of the mountain, fr. 4 Aug. 1974, *Daramola et al.* 71111 (K barcode K002833088!); Ondo, Idanre Hills; Carter Peak, fr. 29 Oct. 1949, Keay 25503 (K barcode K002833086!); Ondo, Idanre Hills, on east side of Orosun, fl. & fr. 30 Oct. 1949, Keay 25512 (K barcode K002833087!); Idanre hills, 30 km E of Ondo, about 3 km S of Idanre fl. & fr. 1 Oct. 1977 (K barcode K002833089!)

##### Conservation status

This species is only confidently known from the Idanre Hills in Ondo province, western Nigeria. This small area of granite inselbergs only covers a few square kilometres, such that the extent of occurrence is inferred to fall within the threshold for the Critically Endangered category (< 100 km^2^). It has only been collected from two of the inselberg ‘hills’, although it is likely that it occurs on additional, less accessible outcrops which have not been thoroughly surveyed. However, the total area of potentially suitable habitat at Idanre Hills is small; Richards (1957) observed that the *Trilepis* (now *Afrotrilepis*) mat community which *H. fruticosa* occurs in, covered only about 20% of the surface of Carter’s Peak, one of the two inselbergs which it has been recorded from. Thus, the area of occupancy is likely to be <10 km^2^. There is inferred to be a continuing decline in the area, extent, and quality of habitat for this species due to wood cutting and clearance for cultivation of cash crops such as cocoa, pepper, corn, and bananas (Onadeko *et al*. 2014). Given the extremely restricted range and continuing decline of habitat, this species is preliminarily assessed as Critically Endangered B1ab(iii)+B2ab(iii).

##### Phenology

Collected in flower and fruit in Oct.

##### Etymology

Named for the shrubby growth habit which distinguishes it from co-occurring species of *Heterotis*.

#### 4. Heterotis buettneriana *(Cogn. ex Büttner) Jacq.-Fél.* (Jacques-Félix 1981: 418)

Lectotype (first-step lectotypification designated by Jacques-Félix (1971); second step-lectotypification designated here): Gabon, Ogôoué, *Thollon 444* (P barcode P00412569! image; isolectotypes: P00412568! image, P00412570! image)

Basionym*: Osbeckia buettneriana* Cogn. ex Büttner (Büttner 1889: 95; Cogniaux 1891:332)

*Dissotis buettneriana* (Cogn. ex Büttner) Jacq.-Fél. (Jacques-Félix 1971: 547)

*Osbeckia rubropilosa* De Wild. (De Wildeman 1922: 378) Type: Democratic Republic of the Congo, Avakubi, 30 Dec. 1913, Bequaert 1678 (BR! image)

##### Distribution and habitat

Cameroon, Equatorial Guinea Gabon, Republic of Congo, Democratic Republic of Congo. Lowland evergreen forest and clearings. 50 – 800 m elev.

##### Representative specimens examined

**CAMEROON.** Central Province, Mefou Proposed National Park Ndanan 2, 03.37°N 11.34° E, 700 m, fl. & fr. 22 March 2004, *Darbyshire et al. 185* (K barcode K000023198!, YA, P); Central Province, Mefou Proposed National Park Ndanan 2, 03.37°N 11.34° E, 700 m, fl. 15 Oct. 2002, *Cheek et al. 11111* (K barcode K000023197!, YA, WAG, SCA). **EQUATORIAL GUINEA.** Bata-Senge: Estrada km. 38 Bibogo, 650 m, fl. & fr. 9 April 1997, *Carvalho 6279* (NY! image, US! image, WAG! image, MA). **GABON**. Woleu-Ntem, chantier Oveng vers Mitzic, 0.40°N, 11.25 E, fl. & fr. 6 May 1986, *Louis 2163* (K barcode K002834366!, WAG); Ogooue-Maritime, former extraction road system accessible from Peni CBG chantier, 2°02.50’ S, 10°24.98’ E, fl. 28 Oct. 2005, *van Valkenburg et al. 3013* (K barcode K002834367!, WAG). Estuaire, Libreville. P1: Arboretum de Sibang, 0° 25’ N 9° 29’ E, 50 m, fl. & fr. 7 Dec. 1999, *Simons et al. 291* (WAG! image, MO! image, HNG, LBV). **REPUBLIC OF CONGO.** Goumina, (Mayombe), 100-200 m, fl. 1 dec. 1990, *la Croix 4986* (MO! image, BR! image). **DEMOCRATIC REPUBLIC OF CONGO**. Haut-Zaire. Ituri; Zone de Mambasa; ca. 14 km SSW of Epulu and 2 km S of Tito River on trail to Ituri River, summit of Mabobo Hill, 01° 16’ 23"N 28° 32’

27"E, 790 m, fl. 2 Feb. 1994, *Gereau et al. 5273* (BR! image, MO! image, WAG! image); Ituri; Zone de Mambasa; Basakwe Camp, west bank of Basakwe River ca. 4 km ESE of confluence of Edoro and Afarama Rivers, 01°32’24"N 028°32’41"E, 790 m, fl. 10 Feb. 1994, *Gereau & Bocian 5314* (MO! image); near Bafwasende, fl. & fr. 26 Sept. 1957, *Croockewit 700* (WAG! image).

##### Conservation status

This species has a large extent of occurrence (EOO) of at least 900,000 km^2^. It has been recorded from many more than ten locations, occurs in a range of different habitats, including disturbed sites. It is not thought that there any major threats impacting this species. It is preliminarily assessed as Least Concern.

##### Phenology

Collected in flower and fruit between Sept. – April.

##### Etymology

Named for the German botanical collector Richard Büttner.

##### Notes

In the absence of flowering material with which to examine the stamens, this species may be confused with *H. prostrata* due to its distinctly excurrent emergences at the apex of the sepals, which persist in fruit.

The protologue for this species’ basionym (Cogniaux 1891) lists two collections: ‘Büttner in hb. Berol.’ (*Buettner 23*) and ‘Thollon in hb. Mus. Paris’ (*Thollon 444*). The former was presumably destroyed at Berlin. The extant specimen *Thollon 444* (P) was subsequently designated by Jacques-Félix (1971) as a lectotype, however there are three sheets for *Thollon 444* at P. We designate the *Thollon 444* sheet with barcode P00412569 as the second step lectotype because it is the only sheet with flowers in which the characteristic isomorphic stamens are clearly visible (Art. 9.17 of Turland et al. 2018).

#### 5. Heterotis decumbens *(P.Beauv.) Jacq.-Fél*

(Jacques-Félix 1981: 418) Type: *Palisot De Beauvois s.n.* (G G00014591! image)

Basionym*: Melastoma decumbens* P.Beauv (Palisot De Beauvois 1806: 69)

*Rhexia decumbens* Poir. (Poiret 1816: 628)

*Osbeckia decumbens* DC. (Candolle 1828: 143)

*Asterostoma decumbens* Blume (1849: 50)

*Dissotis decumbens* Triana (1872: 58)

*Heterotis laevis* Benth. (Bentham 1849: 348) Type: Nun River, *Vogel 36* (K barcode K000313165!)

*Dissotis laevis* Hook.f. (Hooker 1871: 451)

*Dissotis mahonii* Hook.f. (Hooker 1903: 7896) Type: coll. Uganda 1901, cult. Kew 8 Sept. 1902, *Mahon 28* (K barcode K000313072!)

*Dissotis decumbens* Triana *var. minor* Cogn. (Cogniaux 1891: 369) Type: Democratic Republic of Congo, Bangala, fl. & fr. 8 June 1888, *Hens 164* (K barcode 000313104!, BR BR0000006493417! image, BR0000006494384! image)

##### Distribution and habitat

Nigeria, Cameroon, Equatorial Guinea, Gabon, Republic of Congo, Democratic Republic of Congo, Central African Republic, South Sudan, Uganda, Tanzania, Angola. Introduced to Madagascar and the Mascarene Islands. Grows in forests and forest margins or clearings, along streams, roadsides, in plantations, and disturbed areas, sometimes creeping on boulders. 0 – 1, 200 m elev.

##### Representative specimens examined

**NIGERIA.** Degema, Brass, fl. 14 December 1962, *Daramola & Aderson FHI 46329* (K barcode K002834257!) **CAMEROON.** Abong-Mbong, fl. 10 Sept. 1962, *Price & Evans 254* (K barcode K002834403!); South Region, south of Kribi, near new Kribi deep water port site, 2°42’18.0"N 9°51’49.1"E, fl. & fr. 16 October 2017, *van der Burgt et. al. 2151* (K barcode K001286879!, WAG, YA). **EQUATORIAL GUINEA.** Fernando Poo, fl. 27 Nov. 1946, *Guinea 258* (K barcode K002834410!).

**GABON**. Estuaire, between Cap Santa Clara and Cap Esterias, 0°32.86’ N, 9°18.51’ E, 2 m elev., fr. 23 Feb. 2003, *Wieringa 4812* (K barcode K002834405!, WAG); Ogooue-Maritime, old logging road leading southward from chantier CBG Peni, 2°07.80’ S, 10°25.10’ E, 225 m elev., fl. 22 April 2005, *van Valkenburg et al. 3156* (K barcode K002834406!, WAG); Nyanga, Bame, fishing camp, 60 km S of Mayomba, 27 km N of Congo Brazzaville border, 03°47’ 07’’ S, 11°01’ 01’’ E, fl. 11 May 2001, *Walters et al. 640* (K barcode K002834407!, MO). **REPUBLIC OF CONGO**. Kouilou, Tchimpounga, Zone soleil 1, 4° 30’ 53.7’’ S, 11° 46’ 04.8’’ E, 5 m elev., fl. 5 Dec. 2012, *Mpandzou et al. 1709* (K barcode K001393346!); Kouilov, Point 4, Noumbi, bloc Paris, 4° 5’ 58.0’’ S, 11° 20’ 11.0’’ E, 2 m elev., fl. & fr. 15 April 2013, *Nkondi et al. 652* (K barcode K001393328!). **DEMOCRATIC REPUBLIC OF CONGO.** Boma, fr. 10 Oct. 1931, *Dacremont 68* (K barcode K002834408!, BR, WAG); Jardin D’Eala, fr. 22 May 1946, *Leonard 166* (K barcode K002834424!, BR). **CENTRAL AFRICAN REPUBLIC.** Ndakan, gorilla study area Njéké from M5400 to C 5800, 02°21’N 016°12’E, 350 m elev. fl. 7 May 1988, Harris & Fay 603 (MO! image). **SOUTH SUDAN.** Talanga, 4°01’ N, 32° 45’ E, 950 m elev., fr. 2 Dec. 1980, *Friis & Vollesen 638* (K barcode K002834472!), Khor Atiziri, fl. & fr. 28 Feb. 1870, *Schweinfurth 3093* (K barcode K002834473!). **UGANDA.** Entebe, 3900 ft elev., fl. 1901, *Fin. Michon s.n.* (K barcode K002834555!). **TANZANIA.** East shore, fr. Oct. 1893-1894, *Scott-Elliot 8232* (K barcode K002834558!) **ANGOLA.** fl. 1910 *Gossweiler* 4705 (K barcode K002834745!); near Loango River, fl. Oct. 1921, *Dawe 274* (K barcode K002834743!). **MASCARENE ISLANDS (INTRODUCED). Reunion.** Bord de zoute Le Tremblet, fl. 20 Nov. 1970, *Friedmann 629* (K barcode K002834785!)**; MADAGASCAR (INTRODUCED):** Antsiranana, Fivondronana d’Antalaha, Canton d’Ambohitralanana, à Andrahimbazaha, 15° 17’ S, 50° 27’ E, 0-60 m elev., fr. 29 April 1994, *Rahajasoa 367* (K barcode K002834784!); Toamasina, Soanierana Ivongo, Antanambao-Andrangazaha, 16° 51’12’’ S, 49° 41’37’’ E, 0 m. elev., fr. 27 Nov. 2016, *Rakotoarisoa & Andriamahay SNGF 3840* (K barcode K000936861!, TAN, SNGF, TEF);

##### Conservation status

This species is widespread with a very large range. It occurs at many more than ten locations and is suspected to have a large population size. It is not thought that there any major threats impacting this species. It is therefore preliminarily assessed as Least Concern.

##### Phenology

Collected in flower and in fruit between October – June.

##### Etymology

Named for its decumbent habit.

##### Notes

The stalked, stellate hypanthial emergences which otherwise characterise the genus *Heterotis* are reduced to setae or simple hairs in this species (Fig. 1 H, I). In some specimens, the hypanthium is glabrescent. Whereas *H. prostrata* and *H. rotundifolia* appear to be widely naturalized across the tropics, this species is only known to be naturalized in Madagascar and the Mascarene Islands. All of the specimens of *Heterotis* from Madagascar and the Mascarene Islands examined in the preparation of this synopsis belonged to *H. decumbens*.

#### 6. Heterotis prostrata *(Thonn.) Benth.* (Bentham 1849: 349)

Lectotype (designated here): Ghana *Thonning 285* (C barcode C10004150! image; isolectotype C10004151! image) Basionym: *Melastoma prostratum* Thonn. (Thonning 1827: 220)

*Dissotis prostrata* Hook.f. (Hooker 1871: 452)

*Dissotis rotundifolia var. prostrata* (Thonn.) Jacq.-Fél. (Jacques-Félix 1971: 548)

*Dissotis deistelii* Gilg ex. Engl. (Engler 1921: 748)

*Dissotis schliebenii* Markgr. (Markgraf 1935: 716) Type: Tanzania, Muera Hochfläche, 60 km westlich Lindi, fl. 24 Feb 1935, *Schlieben 6068* (BM isotype! image)

*Lepidanthemum triplinervium* Klotzsch (1861: 64)

*Heterotis triplinerva* (Klotzsch) Triana (1872: 58)

*Osbeckia zanzibariensis* Naudin (1850: 55) Type: *Bojer s.n.* (P P00412583! image)

##### Distribution and habitat

Ghana, Guinea, Sierra Leone, Côte d’Ivoire, Nigeria, Cameroon, Equatorial Guinea, Gabon, Republic of Congo, Democratic Republic of Congo, Uganda, Tanzania, Kenya, Angola, Zambia, Malawi, Zimbabwe, Mozambique. Grows in forests and forest margins or clearings, along streams, roadsides, in plantations, and disturbed areas, sometimes creeping on boulders.

##### Representative specimens examined

**GUINEA:** Kerouane, Langbalema Mt. ridge to N of Kerouane – Bamankoro road 09° 12’ 48’’ N, 9° 9’ 54’’ W, 745 m; fr. 29 Oct 2008; *Darbyshire 561* (K barcode K000615189!, NHGC); **SIERRA LEONE:** fl. 2 April 1951, *Deighton 5389* (K barcode K002834267!); **COTE D’IVOIRE:** Abidjan; market of Adjamé; 3 June 1970, Koning 711 (WAG WAG.1096117! image); **NIGERIA**: Adamawa Division, Vogel Peak area, approx. 84° 25’ N, 11° 50’ E, 920 m elev., fl. 20 Nov 1957, *Hepper 1414* (K barcode K 002834318!); Benue Platue, Jos, Assob escarpment, 09° 30’ N, 08° 40’ E, fr. 22 April 1972, *Wit et al. 1464B* (K barcode K002834320!); South E. State, Ikom, Abu village about 16 miles from Bende/Ayuk village on Abudu road, fl. 1 March 1973, *Latilo 67746* (K barcode K002834325!); **CAMEROON**: South West Province, Kupe-Muaneguba Division, Kupe Village Muanezum trail, 1200 m elev., fr. 28 March 1996, *Etuge 1840* (K barcode K000197738!, YA); South West Province, Mount Cameroon, Idenau, 2 m elev., fr. 17 Oct 1995, Dawson 35 (K barcode K000050228!, SCA, YA); Littoral Region, Ebo Proposed National Park, village, 04° 24’ 22.3’’ N, 10° 9’ 56.9’’ E, 100 m elev., fr. 3 Dec 2019, *Alvarez 34* (K barcode K001310318!); **EQUATORIAL GUINEA**: Bioco, Malabo-Brasilé, km 11, 220 m elev., fr. 25 July 1986, *Carvalho 2090* (K barcode K002834439!); **GABON**: About 20-40 km. NNE of Koumémayong, fr. 13 April 1988, *Breteler, Jongkind, & Dibata 8671* (K barcode K002834427!); **REPUBLIC OF CONGO**: Léopoldville, Thysville, fr. 28 Jan 1950, *Compére 1371* (K barcode K002834458!); **DEMOCRATIC REPUBLIC OF CONGO**: Riviere Kalamu, Makala (ville de Kinshasa), fl. 18 Nov 1976, *Breyne 3486* (WAG barcode WAG.1092457! image); **UGANDA**: Rabongo Forest, approx. 2° 6’ N, 52° E, 1020 m elev., fl. & fr. 12 May 1993, Sheil 1636 (K barcode K002834560!); **TANZANIA**: Magewga Estate. dist. Korogwe, fl. 15 April 1952, *Faulkner 984* (K barcode K002834614!); Magila nr Muhesh, 600 m elev., fl. 8 Nov 1970, *Archbold 1290* (K barcode K002834611!); Amani, fr. 21 Dec 1949, *Verdcourt 8* (K barcode K002834610!); **KENYA:** K7, Kwale District, Gongoni For Res NE side, 30 m elev., fl. 2 June 1990, *Robertson & Luke 6346* (K barcode K002834559!); K7 Kwale District, Shimba Hills, fr. 1 April 1968 *Magogo & Glover 609* (K barcode K002834582); K7 Kilifi District, just north of Mariakani, fl. & fr. 8 March 1977, *Hooper & Townsend 1256* (K barcode K002834580!); **ANGOLA**: Malange, s.d., *Gossweiler 1275* (K barcode K002834782!); **ZAMBIA**: Mwinilunga District. Road to Congo border., 1290 m elev., fl. 14 Nov 1962, *Richards 17208* (K barcode K002834764!); Rapids Mwinilunga, fr. 20 May 1969, *Mutimushi 3229* (K barcode K002834766!); **MALAWI**: Northern Region, Nkhata Bay District. 5 km W of Nkhata Bay, by road in Kandoli Forest Reserve, 550 m elev., fl. 6 March 1982, *Brummitt et al. 16343* (K barcode K002834761!); fl. & fr. 2 Feb 1955, *Jackson 1440* (K barcode K002834758!); **ZIMBABWE**: District Melsetter; Haroni River, 457 m elev., fl. 24 April 1962, *Wild 5733* (K barcode K002834769!); **MOZAMBIQUE**: Moramballa, fr. 30 Dec 1858, *Kirk s.n.* (K barcode K002834752!); Niassa, Cabo Delgado, Palma, andados 14 km de Palma para Nangade, 50 m elev., fl. 18 April 164, *Torre & Paiva 12132* (K barcode K002834747!); Manica, Chimanimani foothills, Maronga, Chiira River, west of Comeni’s compound, 19° 58’ 33’’ S, 33° 05’ 14’’ E, 342 m elev., fl. 14 Nov 2015, *Darbyshire 906* (K barcode K001187553!)

##### Conservation status

This species is widespread with a very large range. It occurs at many more than ten locations and is inferred to have a large population size. It is not thought that there any major threats impacting this species. It is therefore preliminarily assessed as Least Concern.

##### Phenology

Collected in flower and fruit year-round.

##### Etymology

Named for its prostrate (lying flat on the ground) habit.

##### Notes

There are two sheets for *Thonning 285* at C. We designate the sheet with barcode C10004150 as the lectotype, as most representative, following Article 9.12 of the Code (Turland et al. 2018).

Jacques-Félix (1971, 1983) distinguished this species from *H. rotundifolia* based on elliptical rather than broadly ovate to suborbicular leaves, and sparser covering of emergences on the hypanthium, in addition to the character in the key presented in this synopsis: stellate emergences excurrent at the apex of the sepals (not excurrent in *H. rotundifolia*). All of these characters are variable within these two species, which have been synonymized in some past treatments (e.g. Jacques-Félix 1971). We deploy only the excurrent (*H. prostrata*) vs. non-excurrent (*H. rotundifolia*) sepal emergences in the key couplet for distinguishing these species as this appears to be the most consistent and recognizable difference between them. Yet some specimens (*H. prostrata*) exhibit distinctly excurrent stellate emergences whereas in others, it is difficult to determine if the stellate cluster of hairs can be considered slightly excurrent or not. *H. prostrata*, *H. decumbens*, and *H. rotundifolia* may indeed comprise a species complex as suggested by Jacques-Félix (1981, 1995). Alternatively, intermediate forms could represent hybrids between *H. prostrata* and *H. rotundifolia*. Further work is necessary to clarify the species limits in this genus.

#### 7. Heterotis rotundifolia *(Sm.) Jacq.-Fél*

Type: Sierra Leone *Afzelius s.n*. (BM BM000902398! image)

*Kadalia rotundifolia* Raf. (Rafinesque 1838: 101) *Osbeckia rotundifolia* Sm. (Smith 1819) *Asterostoma rotundifolia* Blume (1849: 50)

*Dissotis rotundifolia* (Sm.) Triana (1872: 58)

*Melastoma plumosum* D.Don (1823: 291)

*Heterotis plumosa* Benth. (Bentham 1849: 348)

*Dissotis plumosa* Hook.f. (Hooker 1871: 452)

##### Distribution and habitat

Guinea, Sierra Leone, Liberia, Nigeria, Cameroon, Equatorial Guinea, Gabon, Republic of Congo. Grows in forests and forest margins or clearings, along streams, roadsides, in plantations, and disturbed areas, sometimes creeping on boulders.

##### Representative specimens examined

**GUINEA:** Guinée Maritime, Boké Préfecture, Kabata, Route de Kabat avers l’ouest, 10° 42’ 59’’ N, 14° 34’ 16’’ W, 10 m elev., fr. 24 Nov 2007, *Camara 85* (K barcode K000460542!, NHGC); Simandou Range, Moribadou to Canga East, 08° 35’ 36’’ N, 07° 51’ 28’’ E, 800 m elev., fl. 8 July 2006, *Cheek 13312* (K barcode K000436912!); Préfecture de Coyah, Kakoulima, 09° 46’ 51.7’’ N, 13° 26’ 02.8’’ W, 542 m elev., fl. 1 Dec 2018, *Konomou* 632 (K barcode K001310314!); **SIERRA LEONE**: Gola National Park, central block, 07° 39’ 38.2’’ N, 10° 51’ 34.8’’ W, 370 m elev., fl. 03 Oct 2013, *Sesay 105* (K barcode K001243830!); Northern Region, Tonkolili District, Sula Mts., Bantho Hill ridge above Bongbonga village, WNW of Numbara Hill, 09° 03’ 05’’ N, 11° 40’ 30’’ W, 750 m elev., fr. 02 Dec 2009, *Darbyshire 609* (K barcode K000191460!, SL, FBC); Fourah Bay College, fl. 3 Oct 1967, *Morton & Jarr SL4953* (K barcode K002834268!); **LIBERA**: Zorzor – Gbarnga road, west of St. Paul river, fl. 27 July 1966, *Bos 2168* (K barcode K00283577!); Nimba, 450 m elev., 6 Oct 1971, *Adam 26228* (K barcode K002834278!); 3 miles northeast of Suacoco, Gbarnga, Central Province, fl. 11 Oct 1950, *Daniel 24* (K barcode K002834284!); **NIGERIA**: West, Ekiti, On the new road from Imesi-Igbajo, fl. 23 Oct 1972, *Latilo & Fagemi 67514* (K barcode K002834323!); near Port Harcourt, on cleared ground 7 miles to N.E., fl. 24 Sept 1942, *Taylor 7* (K barcode K002834327!); West state, Ibadan district, bush near Forestry School, fl. 20 Oct 1971, *Wit 734* (K barcode K002834322!); **CAMEROON.** Cameroons Mnt, Above Buea, 1050 m elev., fl. 30 March 1952, *Morton 6733* (K barcode K000050229!); **EQUATORIAL GUINEA**: Canetera de Musola, Fernando Poo, fr. 11 Jan 1947, *Guinea 1259* (K barcode K002834440!); **GABON**: Ogooué-Maritime, Rabi-Kounga, c. 01° 42’ S, 09° 52’ E, fr, 16 Nov 1991, *Schoenmaker 152* (K barcode K002834444!); Mitzic, 00° 47’ N, 11° 34’ E, 800 m elev., fl. 28 Aug 1957, *Holderness 282* (K barcode K002834446!); **REPUBLIC OF CONGO**: Lekoumou Préfecture, Yakatopema (Moukouma), New MPD Camp. Forest patch at side of old village, Mousaou-junction with road, 500m from new village, 02° 53’ 21.0’’ S, 13° 36’ 44.0’’ W, fl. 29 Jan 2009, *Kami 4198* (K barcode K000518748!); fl. 1860, *Grey s.n.* (K barcode K002834454!)

##### Conservation status

This species is widespread with no known major threats. It has been assessed as Least Concern on the IUCN Red List (Ghogue 2020).

##### Phenology

Collected in flower and fruit year-round.

##### Etymology

Named for its round, orbicular leaves.

##### Notes

In the preparation of this synopsis, many specimens determined as *H. rotundifolia* were found to have distinctly excurrent emergences at the apex of the sepals, placing them in *H. prostrata*. In particular, all of the specimens examined from East Africa were either redetermined as *H. prostrata* or exhibited somewhat intermediate morphology. *H. rotundifolia* is likely less widespread in tropical Africa than presumed, possibly restricted to West tropical Africa, as suggested by Jacques-Félix (1971, 1983). Further work is necessary to determine the species limits in this genus and in this treatment we exclude intermediate specimens.

## Discussion

The genus *Heterotis* is widely naturalized across much of the tropics. Most specimens and records from outside of Africa are determined as *H. rotundifolia*, which is classified as a weed or invasive species in several regions, including Puerto Rico, Australia, Singapore, and several Pacific islands (CABI 2014). However, there are also records of *H. prostrata* and *H. decumbens* from outside of mainland Africa, and the distribution of each species outside of their native ranges has not been clarified. In this synopsis, we only list the naturalized occurrences of *H. decumbens* which could be verified with specimens held at K: all of the collections of this genus at K from Madagascar and the Mascarene Islands are *H. decumbens*. Citing Bakhuizen (1946) and Almeda (1990), Jacques-Félix (1994) suggests that early introductions to Java and Hawaii respectively represent *H. rotundifolia*. However, Veranso-Libalah et al. (2017) state that the widely naturalized *Heterotis* across Asia, North America, Oceania, Central America and the Caribbean is in fact *H. prostrata*, wrongly identified as *H. rotundifolia*.

Cheek & Onana (2021) list 25 endemic and 30 near-endemic plant species currently known from Mt Kupe, of which 18 were not yet formally described at the time. The description of these new species is an urgent task; most undescribed species are threatened (Brown *et al*. 2023), yet their conservation status cannot be assessed for the IUCN Red List of Threatened Species until they are formally named. The description and assessment of threatened species is also critical for insuring their conservation *in situ*, for example through the identification and designation of Tropical Important Plant Areas (TIPAs) (Darbyshire *et al*. 2017). We herein contribute to the effort to characterise the endemic and unknown flora of Mt Kupe by formally describing *H. kupensis*, a species which was first recognised as new to science twenty years prior but has remained unnamed until now (Cheek *et al*. 2004). We provisionally assess *H. kupensis* as Vulnerable, providing further support for the protection of Mt Kupe as a critical site for preserving threatened and endemic plant diversity. Mt Kupe is designated as one of Cameroon’s most high ranking Important Plant Areas with 145 threatened species on the IUCN Red List as of 2022, of which 19 are Critically Endangered, within its 145 km^2^ (Murphy et al. 2023). It is not currently formally protected.

The new species described here is best placed in the genus *Heterotis* on the basis of morphological evidence. However, we caution that there is currently no mature fruiting material, precluding the description of seed characters which are an important trait for generic delimitation in Melastomateae, including for *Heterotis*. Further, the staminal morphology is entirely unique for the genus. *H. kupensis* also diverges from *Heterotis* as characterised by Veranso-Libalah *et al*. (2017) in that it is an erect herb with stems that are woody at the base, rather than a decumbent herb. However, we note that both *Heterotis fruticosa* and *H. welwitschii* are also upright subshrubs, and some specimens of typically decumbent or prostrate species in the genus appear to be somewhat fruticose. We therefore suggest that habit is not a reliable diagnostic feature for the genus.

We attempted to generate DNA sequences for the markers used by Veranso-Libalah et al. (2017) in order to provide molecular support for this species’ placement in *Heterotis* but were unable to successfully extract DNA from the type specimen. Future collections of this species may reveal intraspecific variation, seed traits, or molecular evidence which may prompt re-examination of its generic placement within Melastomateae. However, we consider its description an urgent task given its narrow endemicity and apparent vulnerability to extinction, and we therefore describe it here as a morphologically unique member of *Heterotis*.

## Acknowledgements

We thank Felix Forest and Laszlo Csiba for their efforts to obtain marker sequences for *H. kupensis* and Andrés Fonseca-Cortés for his help in making the distribution map for the new species. V.J.S. was funded by the NERC Science and Solutions for a Changing Planet Doctoral Training Programme (grant no. NE/S007415/1), the CASE component of which was funded by St. Andrews Botanic Garden. The botanical surveys in Cameroon which produced the material of *Heterotis kupensis* in this paper were mainly supported by Earthwatch Europe (1993 – 2005) and by the Darwin Initiative of the UK Government through the Plant Conservation of Western Cameroon and the Red Data Book of Cameroon projects, both led by the Royal Botanic Gardens, Kew, working with the IRAD-National Herbarium of Cameroon. Gaston Achoundong, former head of the National Herbarium of Cameroon (YA) and his successors including Jean Michel Onana and Barthelemy Tchiengué are thanked for their collaboration and support over the years. The funders had no role in study design, data collection and analysis, decision to publish or preparation of the manuscript.

